# Monodisperse drops templated by 3D-structured microparticles

**DOI:** 10.1101/2020.03.22.001065

**Authors:** Chueh-Yu Wu, Bao Wang, Joseph de Rutte, Mengxing Ouyang, Alexis Joo, Matthew Jacobs, Kyung Ha, Andrea L. Bertozzi, Dino Di Carlo

## Abstract

The ability to create uniform sub-nanoliter compartments using microfluidic control has enabled new approaches for analysis of single cells and molecules. However, specialized instruments or expertise have been required, slowing the adoption of these cutting-edge applications. Here, we show that 3D-structured microparticles with sculpted surface chemistries template uniformly-sized aqueous drops when simply mixed with two immiscible fluid phases. In contrast to traditional emulsions, particle-templated drops of a controlled volume occupy a minimum in the interfacial energy of the system, such that a stable monodisperse state results with simple and reproducible formation conditions. We describe techniques to manufacture microscale drop-carrier particles and show that emulsions created with these particles prevent molecular exchange, concentrating reactions within the drops, laying a foundation for sensitive compartmentalized assays with minimal instrumentation.

## Introduction

The ability to break up a fluid volume into many uniformly-sized compartments that do not cross-talk underlies a number of applications in life science research and diagnostics. Microfluidic technologies have been used to create uniform isolated volumes in microscale wells^1,2,3,4^, valved chambers^5,6^, or through the generation of monodisperse drops from co-flowing streams of water and oil^7,8,9,10,11^. Breaking up a sample volume into smaller uniform compartments enables the concentration of single entities (e.g. cells or molecules) in a subset of these compartments while minimizing background, leading to increased sensitivity and reduced reaction time. Leveraging these capabilities, microfluidic compartmentalization approaches have led to significant advances in counting individual nucleic acids and proteins (i.e. enabling digital PCR and digital ELISA) ^,12,13,14,15,16^ as well as analyzing individual cells based on their secretions or molecular components. The association of a solid phase with each compartment also enables surface-based reactions and barcoding, which has led to transformative applications in single-cell analysis and chemical synthesis^14,15,17,18,19^, but can be limited by random encapsulation processes^19^.

Although providing significant value, the need for significant microfluidics expertise or new chips and costly commercial instruments to perform compartmentalization and measurement has slowed the adoption of these technologies. In a laboratory setting, expertise in microfabrication and clean room infrastructure is necessary to manufacture microfluidic chips; moreover skills in operation of microfluidic devices and development of custom optical or electronic readers is needed even if one has microfluidic chips available. Alternatively, a potential user can acquire commercial instruments that are customized for each particular application (e.g. digital PCR, single-cell RNA-seq, digital ELISA), often with multiple instruments needed to first break up the fluid sample into small volumes, and then analyze those volumes.

A fundamental challenge has been that a collection of droplets in an immiscible fluid are only metastable, requiring energy to create them and surface effects to help stabilize the interface between the two immiscible phases^20,21,22,23^. Precise control of flow rate/pressure with complex instrumentation are needed to stably generate uniform drop volumes, and specialized surfactants are needed to stabilize this out-of-equilibrium state. Coalescence of drops leads to thermodynamic equilibrium, resulting in non-uniform drop sizes that can change with temperature or time. Instead of addressing this challenge by controlling the fluid dynamics of breakup or kinetics of re-coalescence, we focus on engineering the interfacial energy of a drop as a function of volume. We probe how changes to the functional form of this volume-energy landscape could result in the robust creation of uniform drop sizes, thermodynamically promoting drop breakup above a critical volume.

By modulating the volume-energy landscape of a growing drop using microscale particles, we describe a mechanism to create uniform nanoliter-scale aqueous compartments with simple mixing and centrifugation steps. Drops are captured by 3D structured microscale particles – drop-carrier particles (DCPs) – comprising materials with tailored interfacial tensions: an inner hydrophilic layer and outer hydrophobic layer (Fig. 1, Supplementary Fig. 1). We generate uniform drops by mixing, pipetting, or agitating a system with DCPs, aqueous, and immiscible phases. One **drop** is associated with each **particle**, an assembly we refer to as a ****dropicle****, which differs from conventional emulsions that are stabilized by amphiphile surfactants or Pickering emulsions that are stabilized by a multitude of nanoparticles^24,25^. We show how pairs of C-shaped DCPs moved apart in space split volumes unevenly above a critical volume, with one DCP associated with a preferred volume. Multiple such pairwise interactions can give rise to a strong mode in the distribution of volumes across a set of interacting and splitting dropicles^26^. We provide a framework for understanding and controlling this behavior in terms of engineering the Volume-Energy curve (V-E curve) for a DCP. We also demonstrate an approach to manufacture DCPs at the microscale, overcoming challenges with patterning materials with different wetting properties into a 3D shaped microstructure. Once DCPs are manufactured, tens of thousands of dropicles can be formed simultaneously in parallel by simply pipetting for 30 seconds, which corresponds to kilohertz drop production rates. Finally, we show that drops generated using this approach are compatible with enzymatic bioassays.

**Fig. 1.**
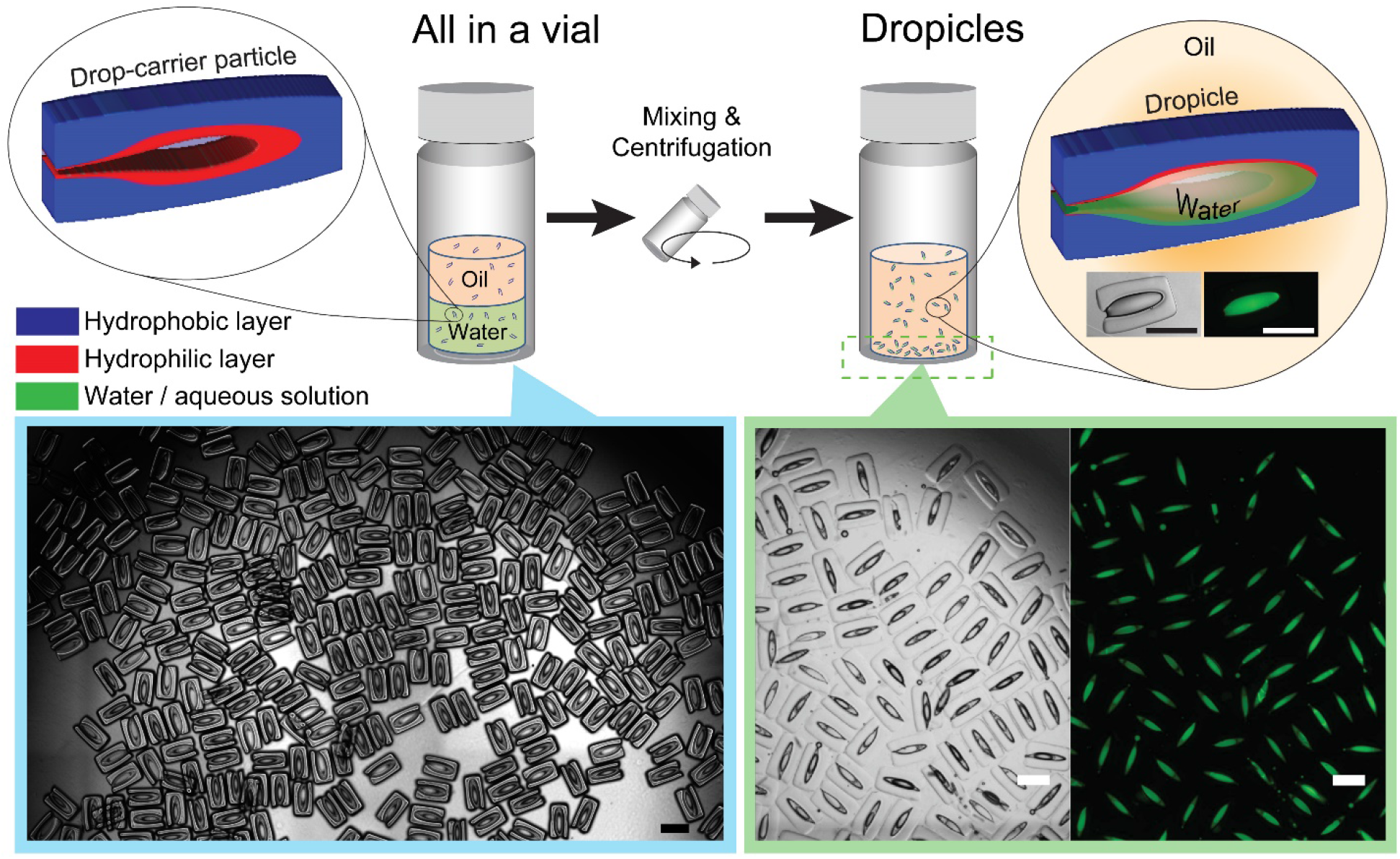
Simultaneous formation of monodisperse dropicles by batch mixing and centrifugation operations. Drop-carrier particles (DCPs) are manufactured with poly(ethylene glycol) and poly(propylene glycol) as the hydrophilic and hydrophobic layers respectively. A collection of DCPs is shown suspended in ethanol on the left. Dropicles with aqueous solution containing fluorescent dye in a toluene continuous phase, shown in the bottom of a vial on the right with brightfield and fluorescence channels. Insets in the right show a single dropicle in brightfield and FITC channels. All scale bars are 500 μm.

A main advantage of our approach is the ability to centrally manufacture drop-carrier particles that can be distributed to end users without expertise in microfluidics and liquid handling. These users can then develop assays using monodisperse nanoliter-scale drops through simple shaking and agitation using widely available laboratory equipment. We expect that a number of assays previously demonstrated using lab on a chip infrastructure could be implemented in this “lab on a particle” format in the future, providing greater access to the deployment of powerful biological assays.

## Results

### Theory of dropicle formation

In a two-phase system, the interfacial energy increases linearly with surface area; for an isolated sphere of volume (V=4πr^3^/3), the energy scales as 4πr^2^ ~ V^(2/3)^ (Fig. 2A), a concave function of volume. For spherical drop emulsions, there is no local minimum in drop size and coalescence of adjacent drops is favored due to the overall decrease in surface area. If the volume vs. interfacial energy (V-E) relationship is instead convex, it is energetically favorable for a drop to split into equal volumes. This process of splitting will continue ad infinitum, again leading to no local minimum in drop size. However, if a V-E curve transitions from convex to concave, a drop splitting into two daughter drops is expected to break evenly for smaller volumes and break symmetry for larger volumes, with one holding a preferred volume close to the inflection point in the V-E curve, and the other containing the remaining volume (Supplementary Fig. 2). For an overall fluid volume exceeding the number of drops multiplied by the preferred drop volume for each drop, this process of asymmetric splitting is expected to accumulate drops with the preferred volume.

**Fig. 2.**
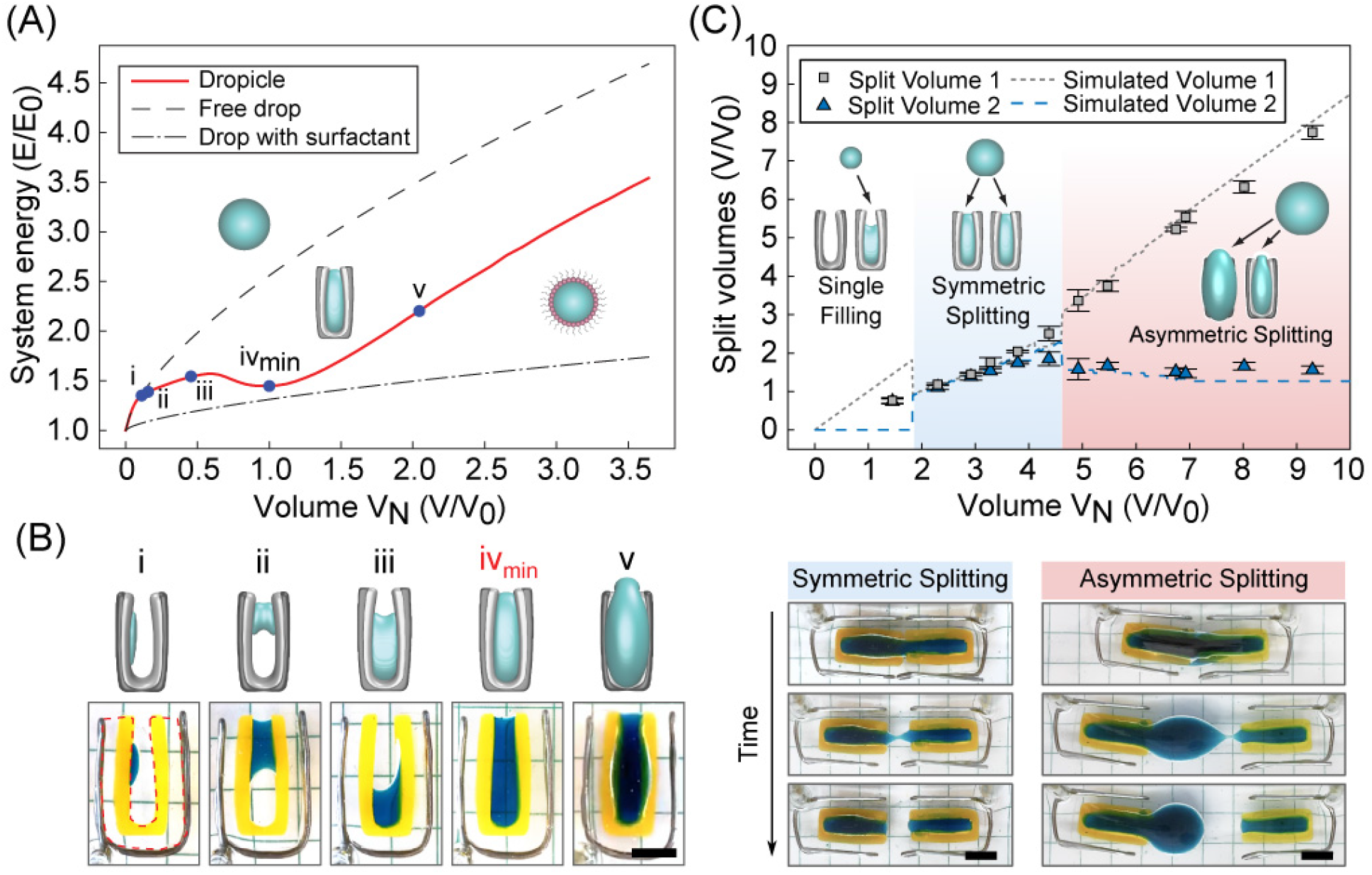
Physics of dropicle generation. (**A**) Simulated Volume - Energy (V-E) curves showing that free drops and surfactant stabilized drops in an immiscible solution possess monotonically increasing energies with V^2/3^, resulting in a thermodynamic driving force for coalescence. Drop-carrier particles (solid red line), however, possess a V-E curve with a local minimum when the drop-carrier particle is substantially filled with fluid (iv). Other configurations of filling are shown along the curve (i-v). The energy and volume corresponding to the local minimum are defined as E_0_ and V_0_. (**B**) Aqueous volumes (blue dye) form different minimal surface shapes when interacting with centimeter-sized DCPs depending on filling volume. The numerical model predictions for (i) to (v) according to (A) and the experimental morphologies of spherical cap, bridge (catenoid), partial filling, and complete filling are shown on the bottom and top respectively. (**C**) Based on the V-E curve in ****A****, splitting of a drop between two drop-carrier particles is theoretically expected to depend on the overall fluid volume (dashed lines), with three regimes of splitting behavior expected. Experimental results (symbols) for centimeter-sized DCPs agree with predictions such that splitting is symmetric within a range of volumes from 2-4, while one daughter droplet is maintained at a preferred split volume above a critical total volume, V_N_ > ~4. Time lapse images are shown for representative experiments in the symmetric and asymmetric splitting regimes.

We hypothesize such a convex-concave functional form is achievable using microstructures at the length scale commensurate with the desired drop size. Practically, an initial concave region of the V-E curve is expected for small volumes as a small drop behaves as a spherical cap on a surface until it achieves dimensions commensurate with the confining microstructure. This “spreading” phase at low volume, in which increasing volume is accompanied by a decreasing rate of increase in surface energy (concave energy), sets the stage for an “inflationary” phase (convex energy) wherein interfacial energy increases more rapidly with increasing volume as the drop fills the microstructure dimensions. Finally, at larger volumes, the V-E curve returns to a concave form consistent with the behavior of a free drop. These conditions are not met with simple topologies such as drops interacting with planes or parallel plates (Supplementary Fig. 3), indicating additional confining surfaces are required, with a trade-off that increasing confinement inhibits drop loading.

### Physical implementation

Drop-carrier particles (DCPs) interacting with a wetting fluid create unique energy minima in the V-E relationship leading to thermodynamic stabilization of drops of specified volumes (Fig. 2). Balancing the need for confined wetting surfaces while also enabling entry of fluid, we design DCPs as C-shaped particles consisting of an inner hydrophilic region and outer hydrophobic layer (Fig. 1). The stable dropicle configuration is simulated using a volume-constrained minimal surface algorithm for the two solid and two fluid phases^16^. The method is an MBO scheme with auction dynamics for the volume constraint (Methods). Our numerical model indicates an initial spreading phase as a low volume of the dispersed fluid forms a single spherical cap (Fig. 2A, location i). A reduced slope in the V-E curve corresponds to the formation of a bridging catenoid (Fig. 2A, locations ii-iii). At intermediate volumes, the drop interacts with more than two surfaces and a local maximum is observed (Fig. 2A, locations iii-iv). Once the interior volume is filled, we observe an inflationary phase in which energy increases with volume at an enhanced rate (Fig. 2A, between locations iv-v). At even larger volumes, the behavior approaches the asymptotic condition of a spherical drop (Fig. 2A), returning to a concave V-E relation. Therefore, DCPs interacting with a fluid volume yield V-E curves satisfying sufficient criteria to split asymmetrically and would accumulate preferred volumes based on our theory (Fig. 2B).

The model also provides information on the contact angles that support stable drops for this DCP design. Generally, the dispersed phase should wet the internal region of the DCP (θ_in_<90) and the external region should not be more wetting than the internal region (θ_out_> θ_in_). Outside of this regime, complex non-filling configurations were observed in our simulations. Additional considerations for practical design of DCPs are also necessary (Methods).

### Experimental observation of asymmetric splitting

We experimentally observe splitting behavior for a volume spanning two centimeter-scale DCPs that are slowly separated. The system is large enough to precisely control the position of neighboring particles while small enough so that capillary effects dominate the mechanics (Fig. 2B-C, Video S1-S2, Supplementary Fig. 4). We adjusted the density of the fluids and separation speed of DCPs to maintain a Bond number and Reynolds number << 1. For example, we utilized PPG as a continuous phase to match the density between the aqueous and oil phases. At different aqueous volumes in a single DCP, we experimentally observe transitions in drop morphology matching the theoretical transitions from a spherical cap, to bridging catenoid on the narrowest approach of the C-shape, followed by filling of the inner cup of the C, and finally wetting of the entire inner surface and filling of the interior volume (Fig. 2B). For the splitting of drops spanning two DCPs, we observed two main regimes that strikingly matched theoretical predictions based on the V-E curves (Fig. 2C). Instead of splitting evenly for all volumes, we found there was one regime where daughter volumes were partitioned evenly (total volumes, *V_N_*, of ~ 2-4*V_0_*, where *V_0_* is the volume at the local minimum of energy), but above a critical total volume of ~ 4*V_0_* one of the daughter volumes is at a fixed preferred volume, independent of the total volume. For example, for total volumes > 4*V_0_*, the smaller daughter drop maintained a quite uniform preferred volume of 1.59±0.14*V_0_*. Notably, the volume with energy minimum at *V_0_* falls close to the inflection point volume of 2.09, where we see a change in curvature from convex to concave in the V-E relation for a DCP. This analysis can be extended to multiple DCPs holding a range of different fluid volumes that are merging and splitting as they are mixed. Given the asymmetric splitting behavior this system is expected to lead to accumulation of DCPs holding the preferred volume while one DCP holds any remaining volume.

### Optical transient liquid molding enables manufacture of microscale drop-carrier particles

We manufacture DCPs at two orders of magnitude smaller length scale (~ 100 μm), addressing the challenges of manufacturing particles (i) comprising two materials with differing interfacial energies, viscosity, and density in complex shapes at the microscale, and (ii) scaling the manufacture to automatically produce a sufficient number of uniform particles for large scale experiments. We manufacture DCPs using an optofluidic technique we developed called optical transient liquid molding (OTLM)^27^, in which we co-flow separate pre-polymer solutions of poly(ethylene glycol) diacrylate (PEGDA) and poly(propylene glycol) diacrylate (PPGDA), shape the streams to the desired cross-sectional morphology in a microchannel flow, and then photo-crosslink this configuration. The particle shape is sculpted along one direction using inertial fluid effects and in an orthogonal direction using photolithographic processes^27,28^ (Fig. 3A, Methods).

**Fig. 3.**
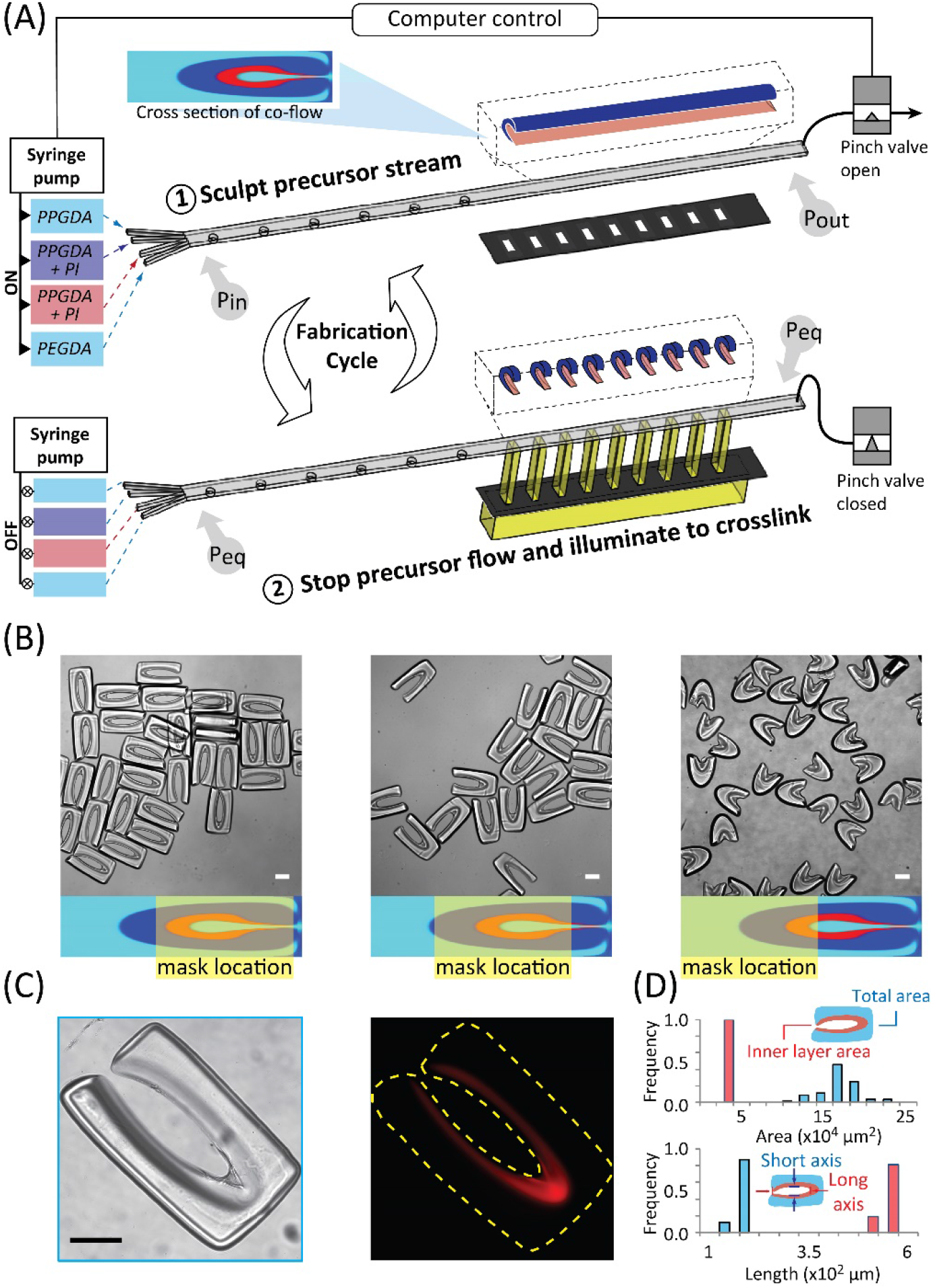
Manufacturing of microscale drop-carrier particles with uniform dimensions. (**A**) Polymer precursors of poly (ethylene glycol) diacrylate (PEGDA) and poly (propylene glycol) diacrylate (PPGDA) are co-flowed with and without photoinitiator. The co-flowing streams are shaped using inertial flow sculpting to create a concentric C-shaped structure in the cross-section of the flow with PEGDA internal to PPGDA. A pinch valve is then closed to stop the flow leading to pressure at the inlet (P_in_) and outlet (P_out_) to equalize to an equilibrium pressure, P_eq_. The sculpted stream is exposed to ultraviolet (UV) light through a mask to polymerize the PEGDA and PPGDA regions mixed with photoinitiator (PI). The valve is opened and polymerized particles are collected before the cycle is repeated. (**B**) Images of three types of DCPs manufactured by shifting the patterned UV mask along a direction perpendicular to the precursor flow. Enclosed DCPs, shown on the left, are used for most of the studies in this work. (**C**) Brightfield and fluorescent images of a DCP after incubation with resorufin, a red fluorescent molecule which partitions into the inner PEG layer. (D) DCP dimensions are reported for a batch of 90 particles, showing the uniformity of the manufacturing process. Scale bars are 100 μm.

We successfully manufacture microparticles comprising two separate materials with different miscibility properties, and substantial differences in viscosity^27^. When two precursor fluids are employed in OTLM, the precursors should be miscible with each other to avoid the effects of finite interfacial tension at the interface of the co-flow which would act against the deformation generated by flow inertia. Here, we leveraged the ability to make PEGDA and PPGDA miscible with each other when suspended in ethanol. Moreover, to eliminate the asymmetry created by the density difference between the co-flowing streams, which can lead to differential settling over a finite flow stopping time^29^, PEGDA and PPGDA are diluted to 60% and 90% respectively v/v with ethanol so the density of all liquids is matched at 0.987 g/mL. The viscosity of the PPGDA solution (38.9 mPa sec) is approximately five times the viscosity of the PEGDA solution (7.0 mPa sec), however, this difference does not lead to significant changes in the flow shape. We tune the concentration of the photoinitiator (PI) in the two precursors so that the speed of the photocrosslinking is uniform between the two materials in the final cured particles. The concentration of PI is 1.3% and 2.6% in diluted PEGDA and PPGDA respectively. These conditions lead to successful polymerization of both precursors as contiguous particles (Fig. 1, Fig. 3B), where a difference in optical contrast between the two material components of the particles is easily observable. We confirm the presence of the inner PEG layer by incubating with the fluorescent dye, resorufin, which selectively partitions into PEG compared to PPG (Fig. 3C)^30^.

We manufacture DCPs with different shapes in large batches using parallel exposure through a mask aligned along the downstream channel length (Fig. 3A). By shifting the position of the UV illumination through the mask location, different shaped DCPs are formed (Fig. 3B). All three types of DCPs possess an internal PEG region, but differ in the degree of encapsulation of this region by the outer PPG layer. Although all particle types can contain stable aqueous droplets, we focus on DCPs with the highest level of encapsulation (i.e. enclosed DCPs) for most experiments reported herein. Because each exposure yields >30 DCPs using the arrayed mask, we can achieve large batch sizes and throughput of manufacture through multiple cycles of flow shaping, stopping, and UV exposure (where a complete cycle required ~7 seconds). Importantly, the process leads to uniform structures of DCPs across the length of the exposed channel (Fig. 3D). The cost to produce 15,000 DCPs with our current OTLM setup is estimated to be ~$45, which can be theoretically reduced to ~$4 by extending the length of the downstream channel to ~24cm [28].*Particle-shape uniformity.* We measure the dimensions of a population of DCPs in order to assess the reproducibility of the manufacturing process (Fig. 3D). Our theory suggests that the cavity size and wettability of the inner layer of a DCP govern the volume of a dropicle that forms. The internal PEG layer surrounding the cavity had an area of 20,000±1,400 μm^2^ and the short and long axes of the void space encapsulated by the inner PEG layer are 95±9 μm and 451±13 μm. Overall, the dimensions are uniform within 6.57% - a metric that helps define the minimum expected uniformity of the dropicle dimensions.

### Generation of dropicles

The protocol for producing dropicles from DCPs requires no specialized equipment (Methods). We demonstrate successful dropicle formation using a number of continuous phases that are immiscible with water, including poly(dimethylsiloxane-co-diphenylsiloxane) (PSDS), PPG, decanol, and toluene. We identify effective protocols for dropicle generation for several continuous liquid phases. For low viscosity continuous phases like toluene and decanol, we disperse DCPs in the continuous phase first, since interactions with a small volume of aqueous sample is readily achievable through rapid mixing. For high viscosity continuous phases like PSDS, DCPs are first mixed with the aqueous sample to ensure there is enough interaction between the particles and the aqueous phase prior to mixing. We found that dropicle generation is largely insensitive to the process of mixing, e.g., pipetting or centrifugation, and the initial dispersion of DCPs in an aqueous or continuous phase. PSDS is chosen as an oil phase for biological proof-of-concept experiments described herein because we found there was modest transfer of water into this continuous phase over days (47 % and 17% loss in area and fluorescent intensity respectively within 2 days), suggesting compatibility with the timescale of enzymatic reactions (hours) or maintenance of cells (days).

### Monodispersity

Microscale dropicles formed in PSDS and toluene possess a preferred drop volume as suggested by theory and centimeter-scale experiments (Fig. 4A). The microstructure of the surrounding particles not only templates the drops in the emulsion (supporting a nominal diameter, ND, of ~200 μm) but also sustains their shape over a long period of time, resisting the usual coarsening process found in standard spherical drop emulsions. Once created, dropicles in toluene maintained the same mean volume for at least 3 days as long as the dispersed phase was prevented from evaporating (Supplementary Fig. 5) while dropicles formed in PSDS had a slow decay in volume over a 3 day period (Supplementary Fig.6). The drop ND is affected by the total volume of the aqueous phase in the experiment. For volumes less than a saturation value, ~20 fold of the entire void volume of the particles, a high percentage of the population is only partially filled with the aqueous phase (Fig. 4B). Once filled, a strong mode in the distribution of nominal diameter is observed at ~200 μm, in agreement with theoretical predictions that asymmetric splitting occurs above a critical total volume for interacting DCPs, leading to a preferred volume accumulating in daughter drops (Fig. 2C).

**Fig. 4.**
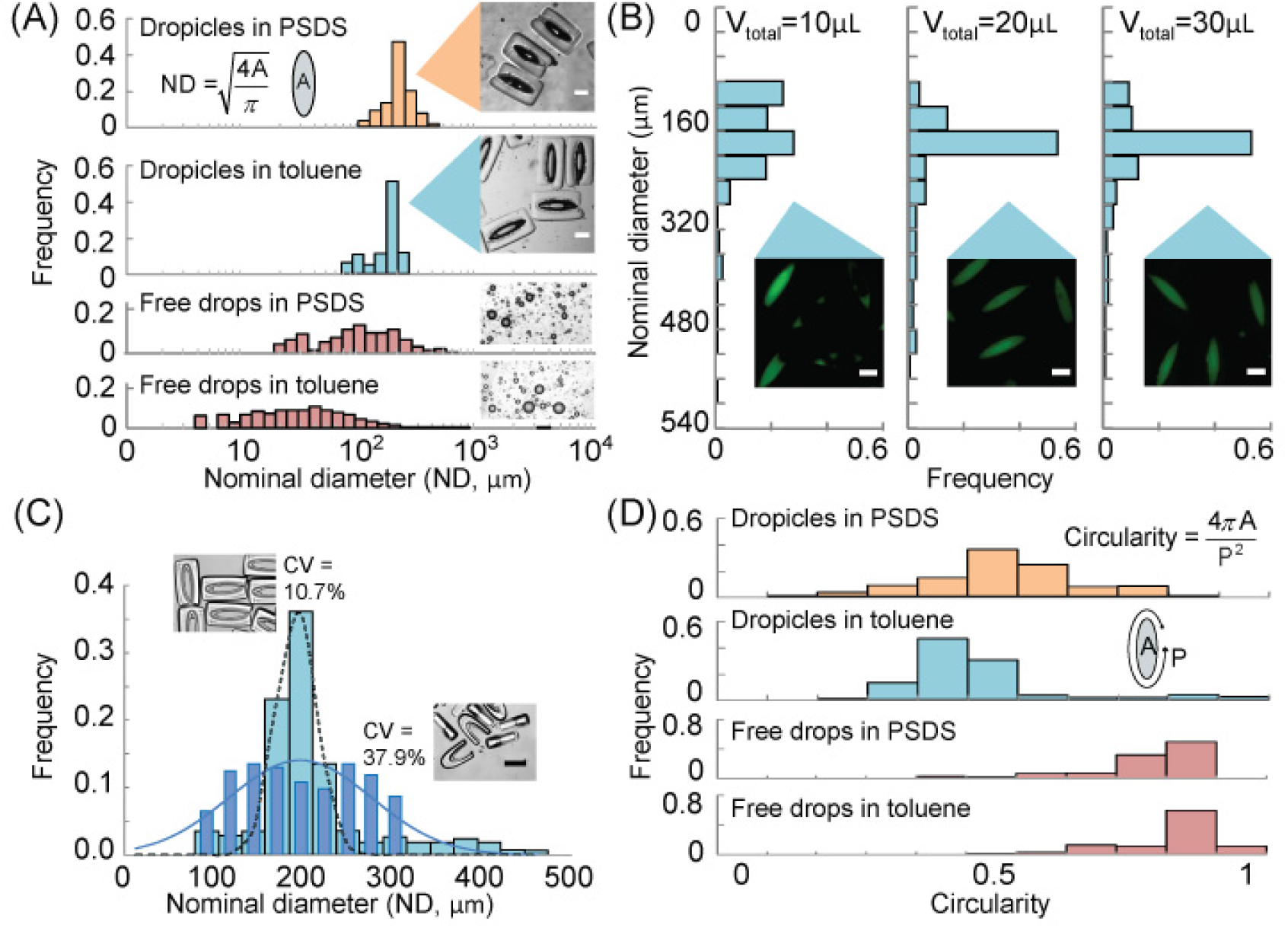
Formation of uniform dropicles. (**A**) Histograms of drop nominal diameters (ND) for dropicles formed in PSDS and toluene. Histograms of free drops in PSDS and toluene stabilized by 0.5% Pluronic surfactant show a much wider distribution in ND (> 100% CV). (**B**) Effect of aqueous volume on dropicle formation. When the volume of the aqueous phase is too low it affects the distribution of nominal diameters among dropicles. The distribution in ND appears to saturate with a mode at 200 μm once the aqueous fluid reaches 20 μL (~20 fold of the total holding volume of the particles). (**C**) Histograms of nominal diameter for enclosed particles and open particles. Increased uniformity is observed for enclosed particles. (**D**) Histograms of a shape metric, i.e., circularity, for dropicles and free drops, showing that dropicles also are stabilized with a unique non-spherical morphology defined by the engineered template. All scale bars are 100 μm.

Drop-carrier particle shape was shown to affect monodispersity (Fig. 4C). Enclosed particles (opening size of 60 μm, which is 6% of the circumference of the interior cavity) had a tighter distribution, CV ~ 11%, and a well-defined mode in droplet ND. However, shorter aspect ratio particles with a wider opening (85 μm, blue diamonds, N=185) have almost four-fold higher variation in size (CV ~ 38%). We observe that two or more particles with the larger opening can stably assemble around a single droplet (Fig. 4C, inset), leading to more variation in drop sizes. Thus a smaller opening is desirable for monodispersity of dropicles.

### Monomorphology

The distribution in the circularity of dropicles formed in toluene and PSDS is shown in Fig. 4D, showing a sharp contrast between a standard emulsion and our engineered system. In agreement with our centimeter-scale experiments, the shape of dropicles minimizes the interfacial energy of the system and is influenced by the DCP cavity shape, whereas surfactant-stabilized drops adopt spherical shapes to minimize energy.

### Dropicles prevent crosstalk

The ability to easily create monodisperse drops supported by a solid-phase opens up many new opportunities for molecular and cellular assays. One fundamental requirement for these assays is the ability to isolate compartments within the system to minimize molecular cross-talk. For conventional surfactant-stabilized droplets, surfactants can potentially enhance transportation of target molecules between phases depending on the properties of surfactants and target molecules [31]. This contrasts with the dropicle system, in which the aqueous droplet is stabilized by only a solid phase without surfactants or with reduced quantity of surfactants, potentially minimizing cross-talk. DCPs also inhibit transfer of dye when agitating dropicles. After mixing dropicles containing separate dye solutions (0.6 kDa and 70 kDa in size), in a toluene continuous phase, we observe the same respective populations of drops without significant exchange of the dye (Fig. 5A, Supplementary Fig. 7). Less than 9.6.% transfer of the 0.6 kDa dye was observed on average while 7.4% transfer of the larger 70 kDa dye was observed after 4 minutes of dynamic agitation by pipetting. This minimal cross-talk following mixing may result from the outer hydrophobic layer of the DCP yielding a physical barrier along with the thermodynamic stability of the supported drops.

**Fig. 5.**
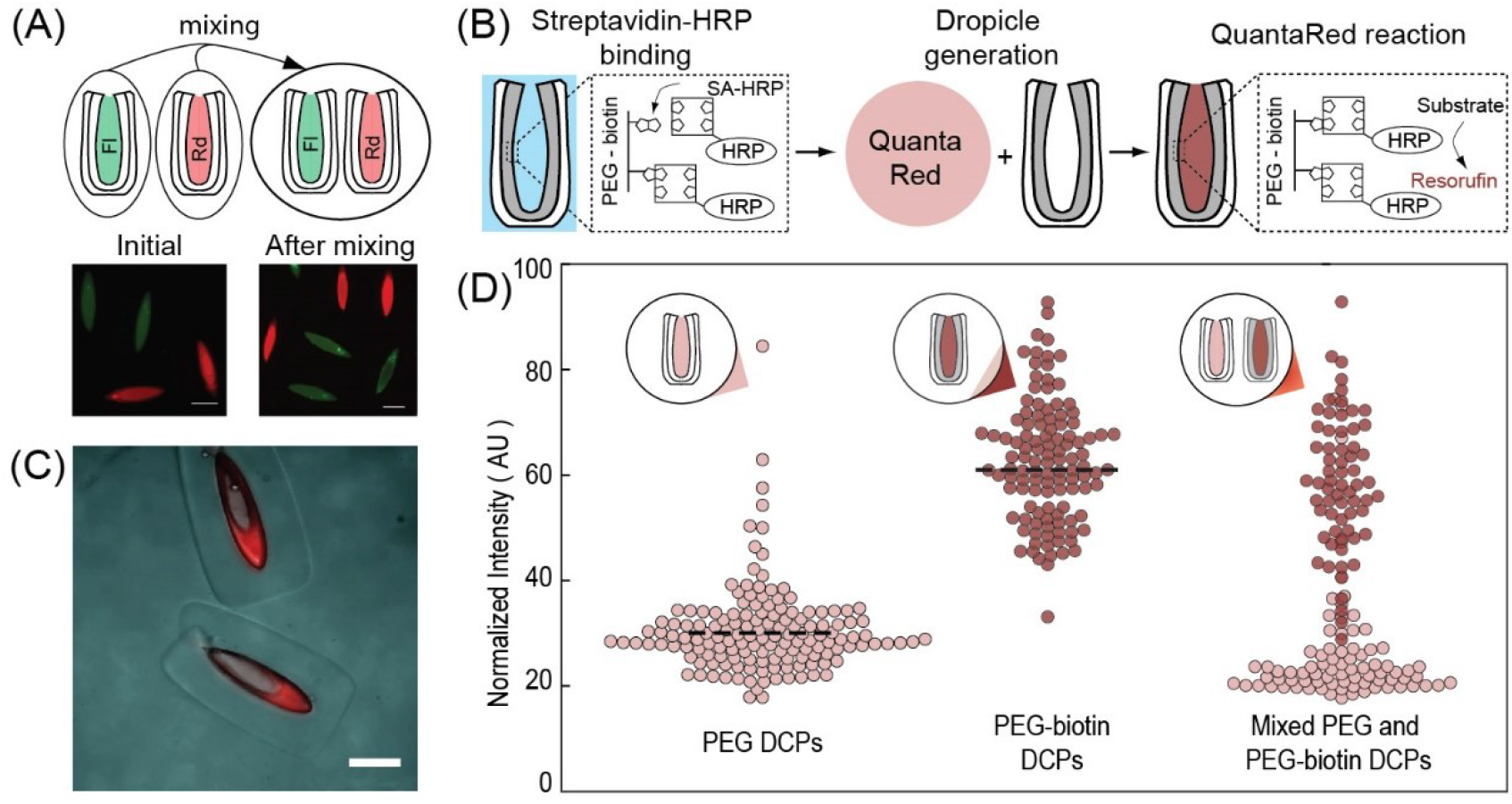
Molecular isolation and enzymatic reactions in dropicles. (**A**) Images taken after introduction and before agitation (left), and after agitation (right) of two groups of dropicles loaded with either biotin-4-fluorescein (Fl, 0.6 kDa, green) or rhodamine B isothiocyanate dextran (Rd, 70 kDa, red) in toluene continuous phase. The dyes do not transfer between dropicles following loading. (**B**) Schematic of the HRP-catalyzed reaction of QuantaRed reagent in which resorufin accumulates within the dropicle. (**C**) Fluorescence images showing the generation of resorufin in dropicles. Fluorescence intensity in the PEG layer increases significantly with the presence of HRP, and is higher than in the surrounding aqueous solution in the dropicle. (**D**) Particles manufactured with and without biotin show selective affinity for streptavidin-HRP and differential generation of resorufin fluorescence. Mixed particles show similar intensity levels to particles in separate wells, indicating minimal cross-talk of generated fluorescent product in a PSDS continuous phase. The dashed lines show the mean for particles with and without affinity to streptavidin. All scale bars are 200 μm.

### Enzymatic assays in dropicles

Leveraging the ability to prevent cross-talk between compartments, we demonstrate a solid-phase enzymatic reaction in which the fluorescent products of the reaction accumulate in dropicles formed in an enzyme-compatible PSDS continuous phase. We modify the inner PEG layer of the DCPs with biotin (Methods), incubate with streptavidin-labeled horseradish peroxidase (HRP), wash away unbound enzyme, and generate dropicles with aqueous QuantaRed reagent (Methods). After generation of dropicles, we incubate the system for various times to generate fluorescent resorufin product. HRP catalyzes the formation of resorufin, which accumulates in a dose-dependent fashion within the dropicle yielding an easily observed fluorescent signal within a 30 minute time period (Fig. 5B-C). Dropicles in which resorufin is enzymatically generated did not cross-talk with neighboring dropicles without reactions. We mix DCPs manufactured with and without biotin with 1 nM streptavidin-HRP and perform the QuantaRed assay as described above. After 24 hours of incubation we can easily distinguish the mean intensity levels in the dropicles with affinity to streptavidin-HRP and those without (Fig. 5D). Notably, the signal for the enzymatic turnover to resorufin shows similar intensity levels in these same particles incubated in separate wells, suggesting transport of product through the oil phase did not contribute to signal intensity (Fig. 5D). Moreover, the resorufin produced also accumulates in the inner PEG layer, yielding a higher fluorescent intensity in this layer, indicating the capability to concentrate signal in this region for future assays. For future applications, non-partitioning dyes can be chosen as the reporter in the assay, i.e., fluorescein. This proof-of-concept suggests that dropicles formed from PPG/PEG layered DCPs can be formed within continuous phases that are enzyme-compatible and prevent cross-talk, two key elements necessary for enzymatically amplified bioassays.

To summarize dropicles can form uniform drops and remain stable while modulating the transport of reporter dyes. Dropicles with a PSDS continuous phase preserve >80% of a small molecule dye after 2 days in static conditions and >90% of 0.6 kDa and 70 kDa dyes after minutes of dynamic agitation. Notably, there is negligible cross-talk over days for resorufin. Exchange in static conditions is likely due to partitioning into the oil phase while in dynamic conditions is likely driven by transient interactions / collisions between dropicles. This is supported by the fact that exchange occurs at almost equal rates for both 0.6 kDa and 70 kDa dyes. Notably, these behaviors contrastwith conventional droplets surrounded by fluorinated oil and formed with surfactant in static conditions. In these conditions transport of fluorescein occurs over days, resorufin over hours, and rhodamine over minutes [32].

## Discussion

There is significant potential, across a range of fields, for the use of thermodynamically stabilized microdroplets associated with solid compartments. The ability for each compartment to be chemically modified with affinity ligands, nucleic acids, or sensing molecules is a key feature for future controlled biological reactions and barcoding. Because each microdroplet is associated with a chemically-defined compartment, and the compartment can be sized to hold only a single particle (Supplementary Fig. 8) or cell, limitations of Poisson loading of cells and beads in standard emulsions can be overcome^19^. Such systems enable single-molecule analysis and synthesis^19,18,30^, or a way to barcode molecules for single-cell analysis^14,15^. The digitized solid structure provides a general substrate to store information from reactions or impart new physical properties into monodisperse emulsions, such as modifications in shape, buoyancy, stiffness, magnetic properties^27^, or stimuli-responsiveness^33^, enabling new opportunities for “lab-on-a-particle” technologies.

## Materials and Methods

### Auction dynamics simulations

#### Droplet Encapsulation Simulation Preparation

We start with a triangulated mesh defining the hydrophobic and hydrophilic surfaces of the drop-carrier particle. This is mapped to a 3D Cartesian grid in which we classify the Cartesian grids into one of four categories: hydrophobic, hydrophilic, droplet, or oil domain. To achieve this, we apply the improved parity algorithm developed in for an Eulerian solvent excluded surface^34^. For a given point x, we draw a half-line emanating from x and count how often it crosses the triangles. The number of crosses determines the phase in which x is located in.

#### Droplet Encapsulation Simulation

In the microscale particle droplet system, the dominant interaction comes from the surface tension between different phases. By ignoring the other forces, we solve for a minimum surface energy configuration using the Auction Dynamics algorithm^35^ on the Cartesian grid. Auction dynamics generates a discrete timestep approximation of volume preserving mean curvature motion of the interfacial boundaries between phases, preserving the volumes of all the phases. As a result, configurations that are stationary under the flow are surface energy minimizers. We iterate the algorithm from an initially spherical droplet on top of the DCP and follow its evolution until it remains stationary under the auction dynamics.

#### Droplet Encapsulation System Post-processing

We compute contact area of each pair of phases to further compute the surface energies of the energy minimization configuration. To systematically address this issue, we first smooth the initial non-smooth sharp interface by running a few steps of Laplacian smoothing. Then we apply the marching cubes algorithm^36^ to extract the level set from the smeared interface. Finally, we triangulate the extracted level set by using the CGAL software and compute its contact area straightforwardly.

### Design considerations for drop-carrier particles (DCPs)

There are additional considerations for practical design of DCPs that are not accounted for in the model explained in the main text. For example, particles should be largely closed such that multi-particle supported drops^37^ are energetically unfavorable and monodispersity is preserved (Fig, 4C). In addition, our model assumes that interfacial energies will dominate the behavior of the system, which is valid when factors such as buoyancy remain small. The Bond number, *Bo* = *Δρgd^2^/Δσ*, for our experimental system is ~ 4×10^−4^, reinforcing this assumption. Here, Δ*ρ* is the density difference between the disperse and continuous phase, *g* is acceleration due to gravity, *d* is the width of the interior void of the drop-carrier particle, and Δ*σ* is the difference between the interfacial tension of the disperse phase and continuous phase with the interfacial tension between the disperse phase and hydrophilic internal material. In our centimeter-scale system we also matched densities to achieve a Bo < 0.1. The interfacial tension of the outer hydrophobic material with the continuous phase should also be small compared to thermal energy to prevent aggregation of particles due to favorable particle-particle contacts on their outer surfaces.

### Microfluidic channel design

We designed the drop-carrier particles using custom software built in lab and open to the public, called uFlow^38^. uFlow enables rapid computation of a 3D particle shape formed from the intersection of an extrusion of the flow stream cross-sectional shape and an extrusion of an orthogonal 2D optical mask shape. Real-time design of the particle shape is possible since the advection maps associated with the inertial flow around a pre-simulated library of pillars is stored and the flow deformation from a pillar sequence is rapidly computed without fluid dynamic simulations. We discovered that six micropillars adjacent to the channel wall can generate a cross-sectional flow pattern with concentric layers with only a small opening on one side, which is suitable for drop-carrier particles when patterned with a rectangular optical mask (see inset of “cross-section of co-flow” in Fig. 3).

### Microfluidic chip fabrication

We fabricate microfluidic chips using soft lithography. The chips contain sequences of pillars designed to create the cross-sectional flow pattern with concentric layers of the precursor materials. The microchannel also contains a long downstream region after the pillars to expose a linear array of patterns to increase fabrication throughput. The silicon mold for replicating poly(dimethylsiloxane) PDMS channels is 300 μm in thickness and thus required a specialized process. We spin a first layer of SU-8 2100 (MicroChem Corp.) to a thickness of 200 μm onto a wafer, recover thermal stress, and spin a second layer of SU-8 with 100 μm thickness. Then, we follow standard protocols for photolithography to develop the mold. We cure PDMS (Sylgard 184, Dow Corning) on top of the mold to replicate the microchannel, peel the PDMS device off the wafer, punch holes for inlets and an outlet, and bond it to a glass slide coated with a thin layer of PDMS using air plasma. The thin PDMS layer matches the surface properties across all walls of the microchannel. The PDMS precursor is spun on the slide at 1000 rpm for 30 seconds and cured in an oven overnight.

### Polymer precursors

Poly(ethylene glycol) diacrylate (PEGDA, M_w_ ≈ 575; 437441, Sigma-Aldrich) and poly(propylene glycol) diacrylate (PPGDA, M_w_ ≈ 800; 455024, Sigma-Aldrich) are chosen to be the polymer precursors for the hydrophilic and hydrophobic layers of the drop-carrier particles respectively. These materials satisfy interfacial tension conditions of importance and are compatible with the OTLM process. The photoinitiator (2-hydroxy-2-methylpropiophenone, Darocur 1173, 405655, Sigma-Aldrich) is introduced with the two precursors.

### Optical transient liquid molding

We us optical transient liquid molding, comprising a flow shaping step followed by a UV exposure step, to manufacture drop-carrier particles^27^. First, we pump a co-flow of polymer precursors into a microchannel with a designed sequence of micropillars at a Reynolds number of 5 to 40. Fluid inertia of the flow around the micropillars leads to an irreversible deformation of an initial rectangular co-flow pattern to a complex cross-sectional pattern. A sequence of micropillars with various sizes and lateral positions can be used to design a wide diversity of cross-sectional patterns, including concave, convex, diamond, stretched bars, etc^39^. Once a pattern is developed downstream of the microchannel containing the micropillars, we rapidly stop the flow and equalize pressure in the channel by simultaneously stopping the upstream pump and occluding the outlet tubing downstream with a pinch valve. Within one second, we illuminate the sculpted precursor stream with a patterned UV light for 500 ms to photocrosslink the precursor stream and solidify multiple 3D-shaped particles. The patterned UV light is created by coupling collimated UV light to a chrome mask with an array of transparent rectangles (140 × 600 μm). Following photocrosslinking, the downstream pinch valve is re-opened and the pump is restarted to flush cured particles into a container outside of the microchannel and to redevelop the precursor flow stream for the next UV illumination cycle. This manufacturing cycle is automated using LabVIEW to fabricate large batches of particles. We also confirm the reproducibility of particle shape across a population of the particles^28^.

After fabrication, all particles are collected in a 50 mL centrifuge tube and rinsed with a volume of ethanol more than 1000 times the sample volume to eliminate the effect of non-crosslinked reagents. The particles were stored in ethanol for later usage.

### Protocol for dropicle generation

#### Protocol for dropicle generation: PSDS

Two approaches are used to generate dropicles using PSDS as a continuous phase. In the first approach we disperse DCPs in an aqueous sample with 0.5% (w/v) Pluronic F-127. We let the DCPs settle in a glass vial and remove the supernatant until the aqueous volume is reduced to ~<50 μL. We inject 1mL of PSDS into the vial and pipette the solution with DCPs, PSDS, and the aqueous phase 1~2 times. The DCPs are left to settle in the vial for about 30 minutes. If needed the supernatant of PSDS is exchanged to remove satellite drops without particles. In the second approach, used to track enzymatic turnover of a fluorogenic substrate in the same particles over time, the particles adhere to the bottom of a well plate for fluid transfer and compartmentalization operations. Specifically, particles suspended in ethanol are transferred to a well plate with a hydrophobic surface (Catalog number: 351143, Corning), and the medium is exchanged after three washes with phosphate-buffered saline (PBS) with 0.5% w/v Pluronic. To characterize the stability of the volume of dropicles over time, food coloring dye (Catalog number: S05189, Fisher Scientific) dissolved in PBS (300 μL) is used to visualize the stability of the volume of dropicles over time. Within seconds, the aqueous solution is fully dispersed around and inside the particle cavity, and excess liquid is removed while an aqueous phase remains trapped within particle cavities. Lastly, 500 μL of PSDS is added on top of the particles to complete the compartmentalization of the aqueous phase.

#### Protocol for dropicle generation: toluene

To reduce the numbers of particle-free satellite drops and adhesion between particles and the glass container, we use a mix of toluene with 10-15% ethanol. Our protocol to create dropicles in toluene differs from PSDS as follows: (1) we disperse drop-carrier particles (initially in ethanol) in 1mL of the toluene/ethanol mix, (2) we inject a small volume of aqueous solution, typically ~20 μL (~17 times of the total void volume of particles), (3) we then pipette the solutions vigorously in a 20 mL scintillation glass vial (VWR) with a hydrophobic coating which is introduced by incubation with Rain-X (ITW Global Brands) for 2 days, (4) following mixing, we centrifuge down the solution in the vial at 2000 rpm for 5 minutes at 25ºC, and (5) and finally pipette away any large visible satellite drops. We cover the vial with parafilm for long-term storage. Moreover, we also confirm that the dropicles can be generated without ethanol using a similar procedure as used for a PSDS continuous phase.

#### Imaging and image processing

We image dropicles and free drops using fluorescence microscopy to evaluate the formation and uniformity in drop size. For clear visualization, 100 μg/mL biotin-4-fluorescein (BF, Catalog number: 50849911, Fisher Scientific) is added to the PBS. We use a custom Python code to analyze the images of the dropicles and free drops. For dropicles, the code detects the fluorescent regions representing drops, filters out regions with size larger than twice or smaller than 0.375 times the nominal size of the particle (corresponding to satellite drops not associated with particles). We measure size/circularity/total intensity for targets, and export an image after filtering, comparing it to the brightfield image for confirmation. For the study of long-term stability, we also filter using circularity to ensure only dropicles are investigated while ignoring spherical satellite drops.

### Method of reaction inside of dropicles

We incorporate additional steps in the protocols for drop-carrier particle manufacture and dropicle generation to perform reactions in dropicles: biotinylation of the inner PEG layer and molecular binding in dropicles. In the fabrication step, we use a mix of PEGDA, ethanol, biotin-PEG-acrylate (Catalog number: PG2-ARBN-5k, NANOCS) in DMSO as the precursor polymer for the inner layer to enable grafting of biotin within the PEG layer during photocrosslinking. After fabrication, we rinse the particles, and store them in ethanol. Prior to use, particles are dispersed in PBS with 0.5% w/v Pluronic, and then incubated with a bulk solution of 1 nM Streptavidin-conjugated HRP (Catalog number: N100, Thermo Fisher Scientific). After binding of the streptavidin-HRP and multiple rinsing steps, we reduce the aqueous volume to ~<50 μL. We mixed ADHP concentrate, enhancer solution, and stable peroxide solution at a ratio of 1:50:50 to make 500 μL QuantaRed solution. We performe reactions within dropicles using two approaches. In the first approach, immediately after the mixing step, we inject the QuantaRed solution into the vial, gently agitated for 5 seconds, inject PSDS, and then pipette the solution up and down to generate dropicles in a glass vial, which took < 2 minutes. The brightfield and fluorescence images of dropicles are taken after 30 minutes of incubation. In the second approach, particles suspended in ethanol are added to a 12 well plate (Catalog number: 351143, Corning). Once particles settle in the well, excess ethanol is removed, followed by three washes with PBS with 0.5% w/v Pluronic. Then, 300 μl of streptavidin-HRP solution at desired concentrations is added and incubated for a given time period, followed by three additional washes. Next, 500 μl of the QuantaRed solution, as described in the first approach, is added to the well to wet the particles, with excess removed immediately. Lastly, 500 μl PSDS is added to form isolated dropicles. Next, fluorescence and bright field images of the dropicles in oil are obtained at desired time points using a fluorescence microscope.

## Author contributions

D.D. conceived of the overall concept of DCPs and dropicles. C.-Y.W. further developed the initial idea, conceived and implemented the fabrication approach to create DCPs, designed protocols to form dropicles, performed experiments, and analyzed data. J.D. and C.-Y.W. performed microgel encapsulation experiments. M.O. developed protocols for dropicle formation and conducted enzymatic amplification and cross-talk experiments. A.J. and J.D. conducted scaled-up experiments and analysis. B.W., M.J. and A.L.B. developed the numerical model and performed modeling of minimal energy configurations. K.H. and A.L.B. developed the analytical framework. All authors contributed to analyzing and interpreting data to formulate the theoretical framework. D.D. wrote the manuscript. C.-Y.W. designed and prepared initial figures. J.D. and D.D. contributed additional figures and modifications. All authors contributed to writing and editing the manuscript and design of the figures. A.L.B. and D.D. supervised the project.

## Competing interests

Authors declare no competing interests.

Correspondence and requests for materials should be addressed to D. D.

---

**Supplementary Fig. 1.**
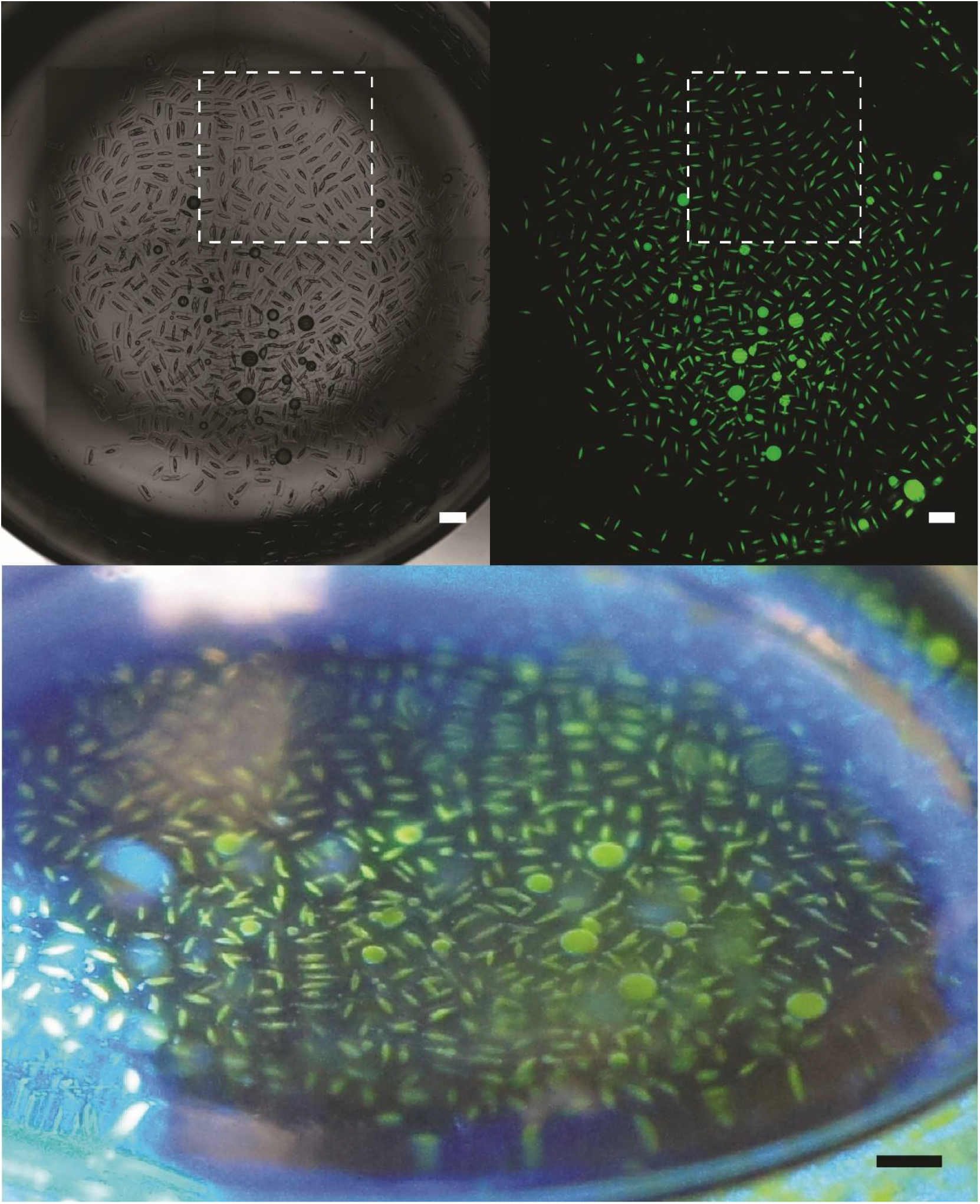
Monodisperse dropicles. Drop-carrier particles, aqueous solution containing FITC-dextran, and oil phases were simply mixed in a scintillation vial and centrifuged down to generate dropicles. The top row insets show stitched brightfield and fluorescence images of the entire vial generated using a microscopy. The white squares outline the areas highlighted in Fig. 1. The bottom image shows an image of the vial from an angle using a standard camera. The top and bottom scale bars are 1 and 2 mm respectively.

**Supplementary Fig. 2.**
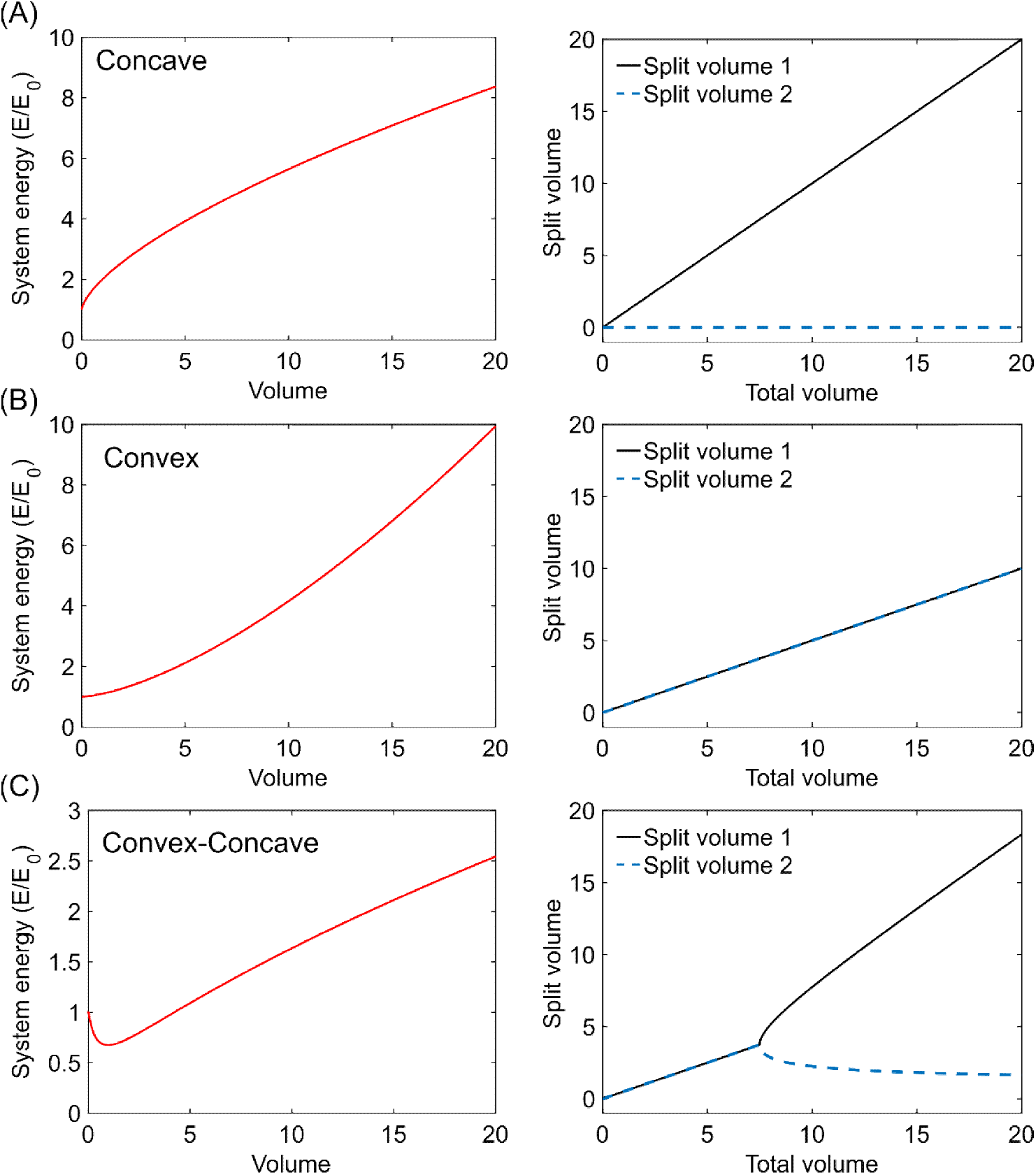
Example volume energy (V-E) curves and corresponding volume splitting plots. (A) For concave V-E curves (e.g. a spherical droplet) it is energetically favorable for volumes to coalesce into a single volume in order to minimize surface area. Equation of the V-E curve: 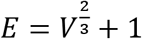 (B) For convex V-E curves it is energetically more favorable for a volume to split into equal volumes. This case also results in no preferred drop volume. Equation of the V-E curve: 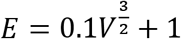 (C) For the case of a V-E curve that transitions from convex to concave, there is an initial volume regime over which droplets split evenly (similar to the purely convex case). However, once the volume reaches twice the volume of the inflection point (V = 2V_I_ = 7.5), volumes split asymmetrically. Here the total volume splits into two volumes, a preferred smaller volume occurring over a large range of total volumes, and a larger volume containing the remaining volume. Equation of the V-E curve: 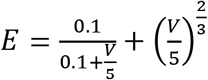

**Supplementary Fig. 3.**
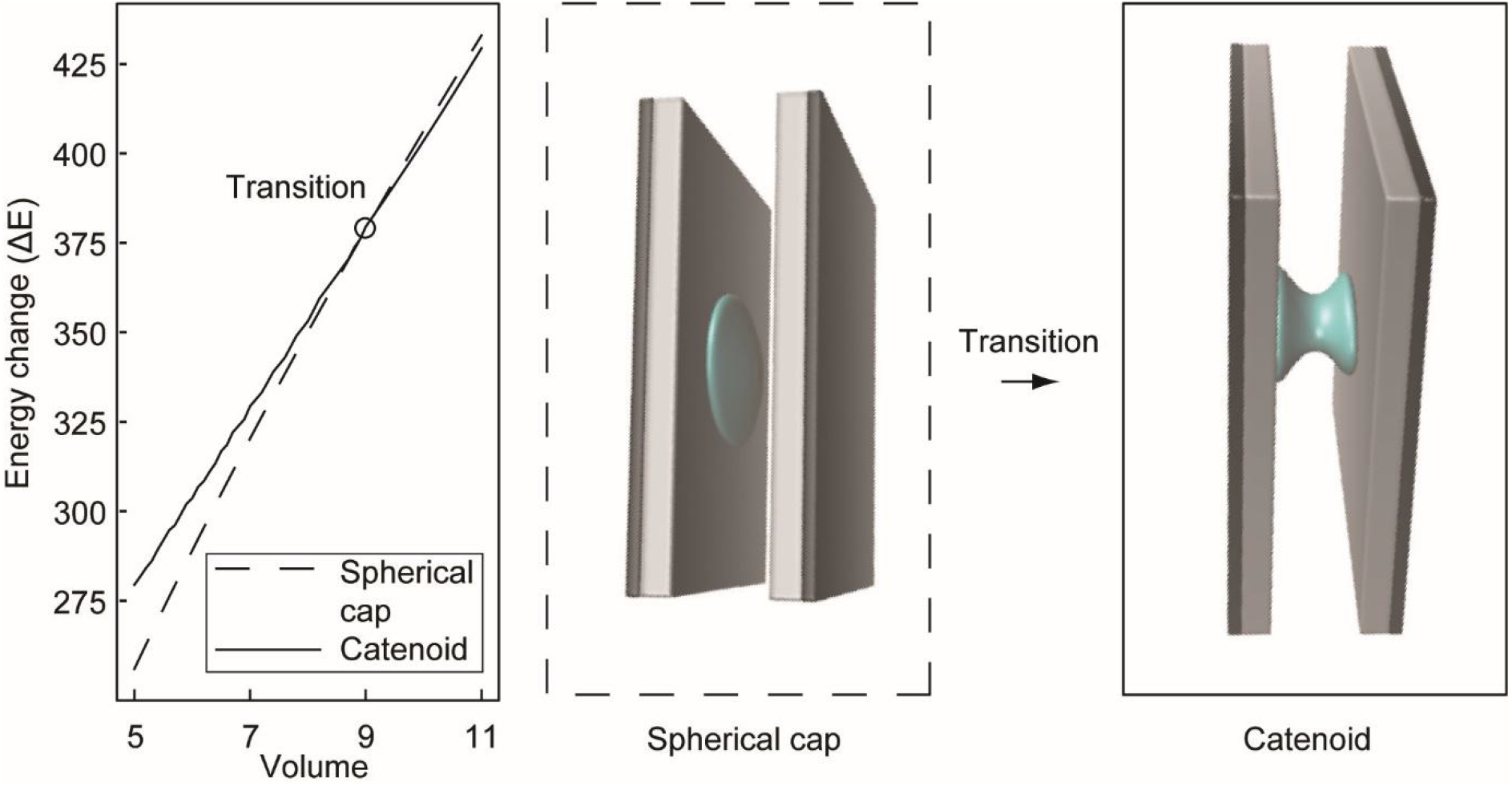
System energy of a drop confined by parallel plates transitioning from a spherical cap to catenoid. The behavior of an aqueous drop with increasing volume is shown using a simplified model of two parallel plates confining the drop with a hydrophilic inner-facing layer. There is a change in morphology of the drop at equilibrium from a spherical cap to a catenoid bridging between the surface, which leads to a change in slope of the V-E curve, however, both remain concave. As shown in Supplementary Fig. 2, additional features in the V-E curve are needed to support monodisperse drops.

**Supplementary Fig. 4.**
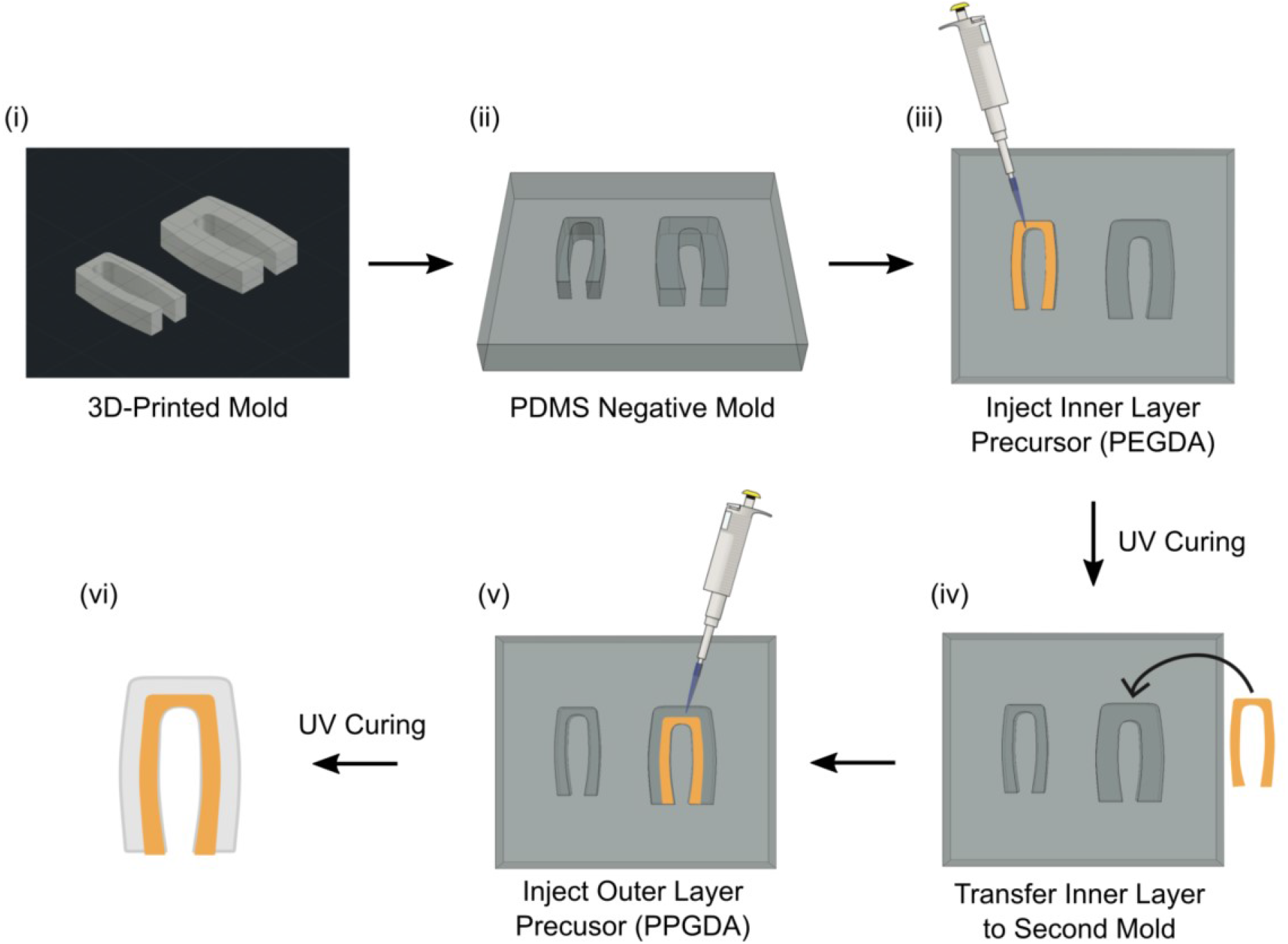
Macro-scale drop-carrier particle (DCP) fabrication process. (i) A positive mold is first printed using an SLA 3D printer (Form 2, Formlabs). (ii) A negative mold is fabricated from the 3D printed mold using PDMS. (iii) PEGDA is pipetted into the smaller mold and crosslinked with UV light (40s, 250 mW/cm^2^) to create the inner layer of the DCP. (iv) The inner layer is transferred into the larger mold, PPGDA is pipetted into the remaining space and crosslinked using UV light (40s, 250 mW/cm^2^). (iv) The resulting amphiphilic DCP is removed from the mold and washed with ethanol prior to volume filling or volume splitting experiments.

**Supplementary Fig. 5.**
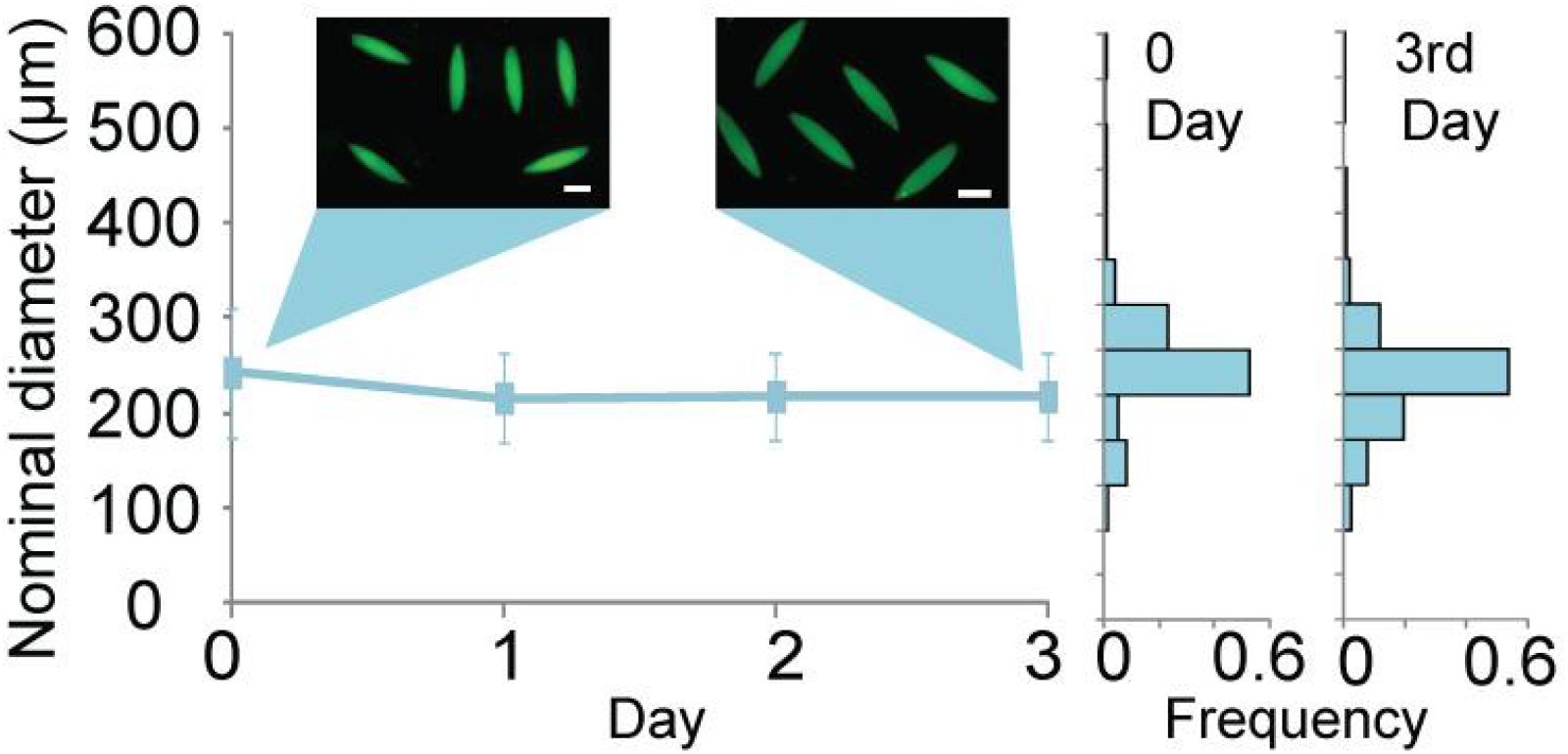
Long term stability of dropicles in toluene continuous phase. The dropicles with fluorescein-containing aqueous solution were generated and imaged on day 0 and day 3. The size distribution of dropicles over three days remains stable. The scale bar is 200 μm.

**Supplementary Fig. 6.**
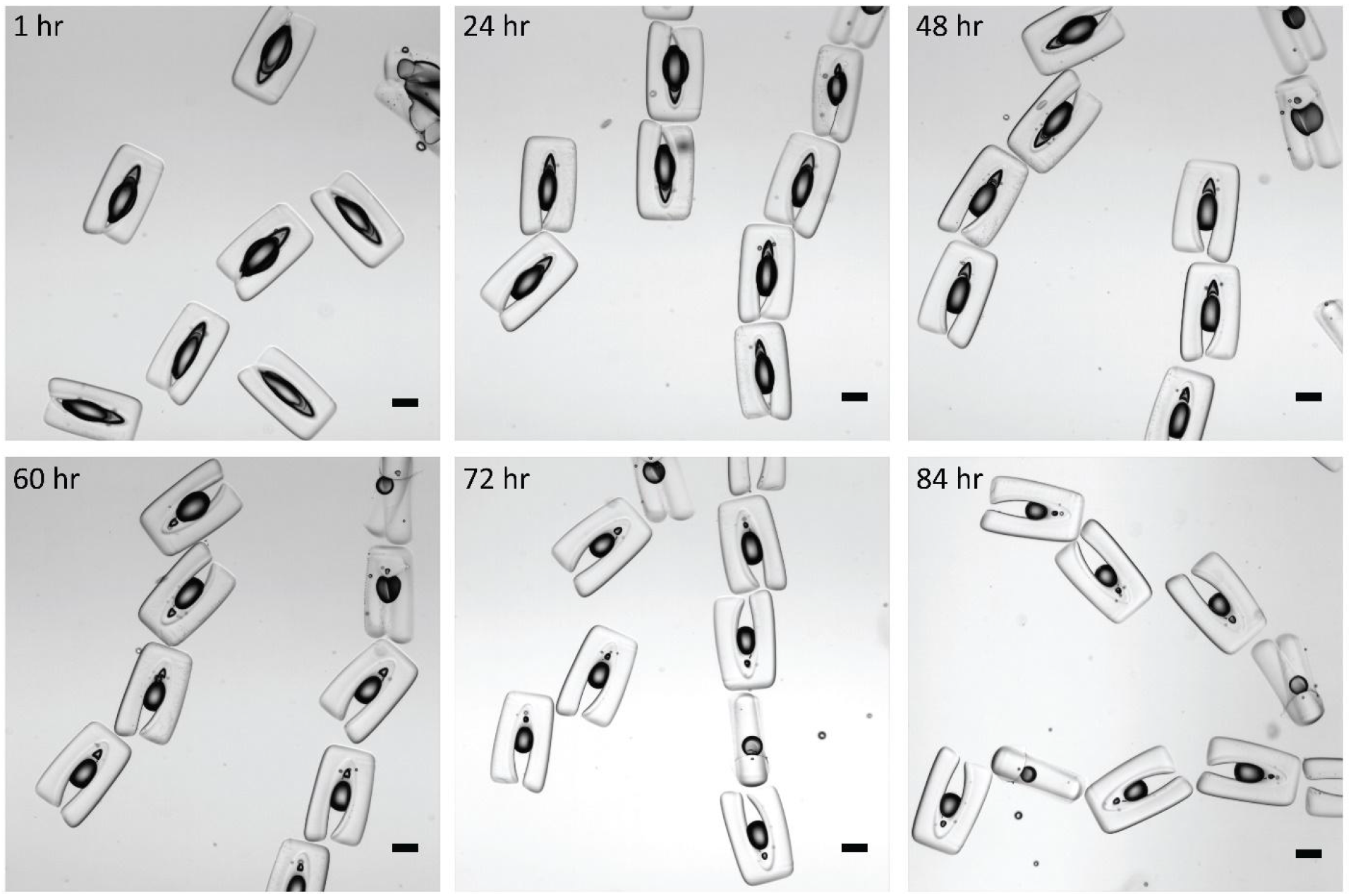
Images of aqueous dropicles formed within PSDS oil are shown over 3 days. The images indicate that the volume can be maintained over several days. A slow reduction in the volume of templated drops is presumably due to dissolution of water in the oil phase and evaporation over time. Scale bar is 200 μm.

**Supplementary Fig. 7.**
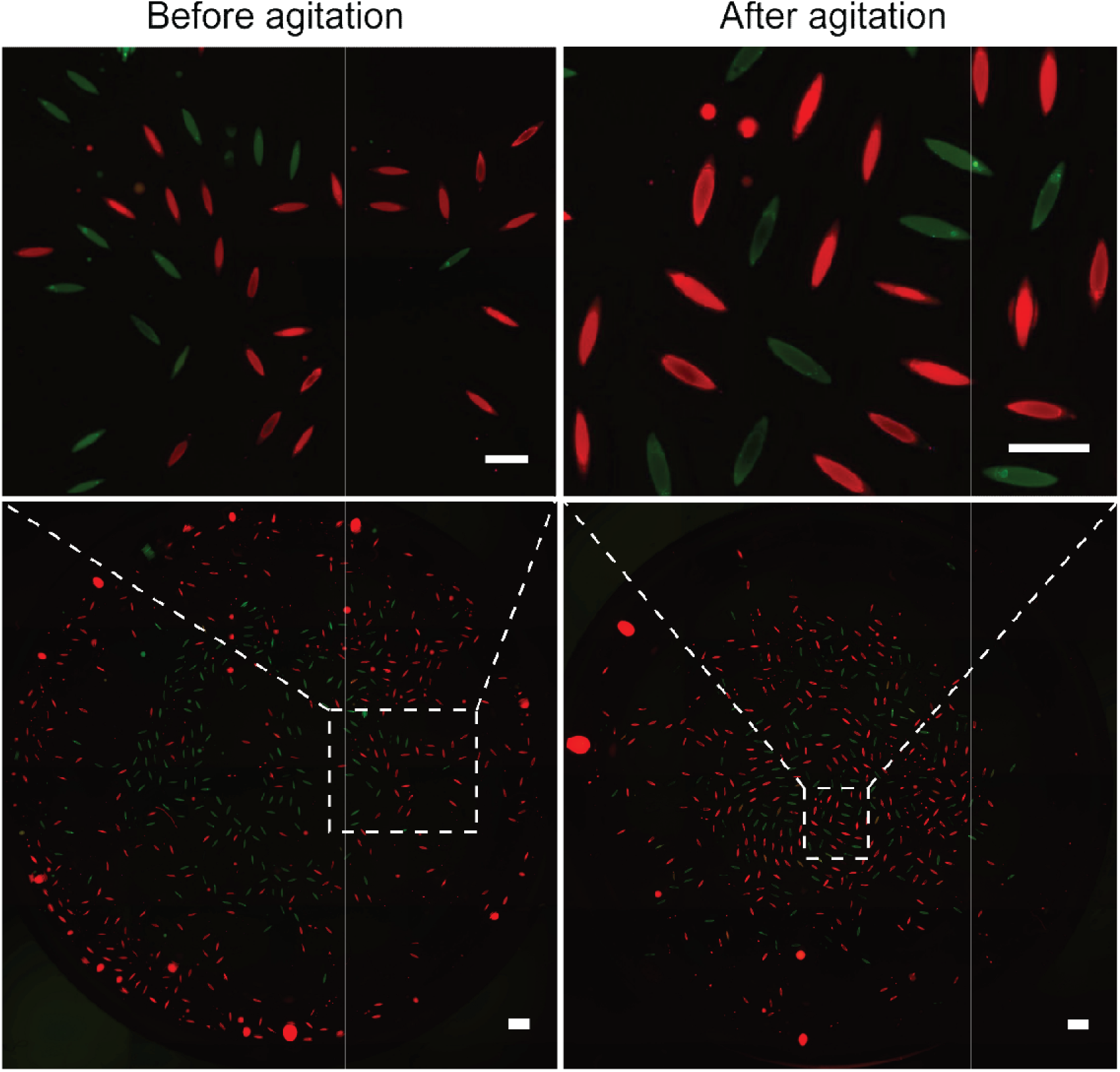
Inhibition of solution exchange between dropicles. We generated dropicles with 10 μg/mL biotin-4-fluorescein (BF) and 1 mg/mL rhodamine B isothiocyanate dextran (RBD) separately in two vials. We introduced ~0.5 mL of dropicle-laden solution from each vial into a new vial. We imaged the blended dropicles in both FITC and TRITC channels before and after shaking the vial on a standard analog shaker (VWR) for 4 minutes. Before and after agitation, only green and red fluorescent drops were observed in the overlay images, indicating there was no transport of dye between solid boundary-protected drops. The scale bars in the top and bottom rows are 500 and 1000 μm respectively.

**Supplementary Fig. 8.**
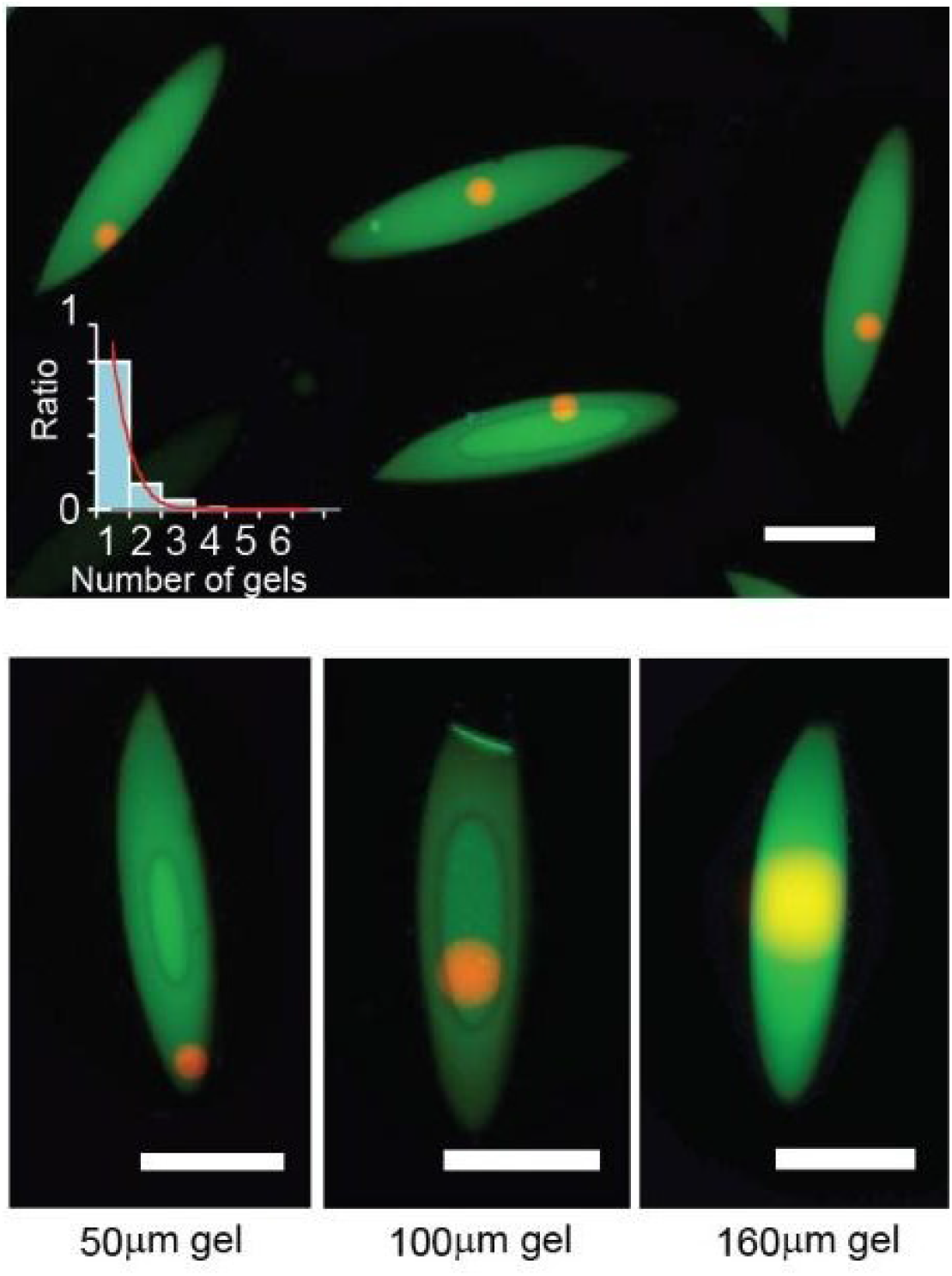
Images of spherical microgels encapsulated in dropicles. The distribution in the number of gels loaded in the dropicles follows a Poisson distribution (inset graph, histogram is experimental results, red line is Poisson distribution). Isolation statistics are independent of size provided the gel is smaller than the drop-carrier particle opening. Microgels are manufactured as described in: de Rutte et al. *Advanced Functional Materials.* (2019): 1900071.

